# HeveaDB: a genetic resource database for rubber tree genomic study

**DOI:** 10.1101/708305

**Authors:** Han Cheng, Yanhua Xia, Jiannan Zhou, Chaorong Tang, Huasun Huang

## Abstract

With the proceedings in the genome study in rubber tree, more and more data were produced. However, these data are still deposited in public databases as raw format, and the utilization of these NGS data is hindered for the researchers in rubber tree community. HeveaDB was therefore constructed and maintained under the organization of International Rubber Research and Development Board. The HeveaDB 1.0 (http://hevea.catas.cn) mainly stores 4 versions of Hevea draft genome, 99 NGS transcriptomes, 18451annotated EST sequences, 12 curated gene families, 30200 gene annotations, 5049 wild germplasm phenotype data, and 18328 Wickham rubber clonal information. Several bioinformatic tools are integrated in the database to facilitate the utilization of the data for the users who are not skilled in bioinformatics. With the periodical update and functional improvement, the HeveaDB will serve as a potential platform for genomic studies and breeding technology in rubber tree.

## Introduction

The Para rubber tree (*Hevea brasiliensis*) is an economically important tropical tree species that produces natural rubber, an essential industrial raw material. Though more than 2000 plant species can produce nature rubber, the rubber trees produce more than 98% of the nature rubber in the world [1]. Now rubber trees are widely planted in the tropical regions of South Asia and Africa, and the produced rubber is one of the major incomes of the small holding farmers in these countries. The *Hevea* genus has about 10 members, all of them are exclusively originated in tropical South America. In 1886, Henry Alexander Wickham collected over 70 000 rubber tree seeds from the central Amazon basin, and send back to the Kew garden, London. About 2300 rubber seedlings was obtained from these seeds. The bulk of seedlings were dispatched to Henarathgoda, Ceylon. Then 22 plants were transported to Singapore in 1877. Thirteen of them went to Kuala Kangsar and nine retained at Singapore Botany Garden. The modern rubber clones are almost all descended from these trees, and they therefore are denoted as Wickham germplasm.

With the proceedings in rubber breeding programs, the yield of rubber tree clones increased substantially. However, due to the long-life cycle, the rubber tree breeding programs takes about thirty years [2], which makes the rubber clone improvement rather difficult. Recent progresses in genomics may accelerate the crop breeding program through the promising genome selection technique [3], in which genome-wide markers are used to predict the breeding value of individuals in a breeding population. However, this technique requires the advances on genetics and genomics. Recent progress in next-generation sequencing technique greatly promoted the genomic study in rubber tree [4–13]. At current stage, 4 versions of draft genome in rubber trees were compiled [14–17] and more than 100 NGS transcriptomic data were produced [18–25]. However, these data are still deposited in NCBI and other public database in raw format, which hindered the NGS data utilization for the scientists in rubber tree community. Besides, huge amount of phenotypic data were produced from rubber tree clones and wild germplasms [26,27]. These data need to be stored and mined in a comprehensive data hub. Considering the present needs in the rubber community, the International Rubber Research and Development Board (IRRDB) decided to construct an integrative genome hub in 2017 International Rubber Conference (Jakarta), which would be used to store the *Hevea* draft genomes, germplasm genome re-sequencing data, clone genome variation data, transcriptomic data, proteomic data, and phenotype data *etc*., and to provide a platform for data archiving, distribution and utilization in rubber tree community. The HeveaDB 1.0 database (http://hevea.catas.cn) was therefore constructed and provided online data access.

## Methods

### Genome data collection and annotation

Basically, two types of rubber tree data were deposited in the HeveaDB: nucleic acid and clonal information data. So far, 4 versions of genome drafts were published in rubber tree, among which the CATAS7-33-97 genome has less scaffold number and longer N50 [28]. Totally 30 200 functional genes were annotated from 43 791 predicted genes. The HeveaDB uses the CATAS7-33-97 genome as the reference. Totally 43 791 protein coding genes were predicted from the reference genome. The expression data were obtained from the published transcriptomes. Totally there are 142 transcriptomes in the GenBank SRA database, among which 99 high quality samples were selected (> 1 Gb) for gene expression calculation. Totally 18451 EST sequences were collected, in which 15 870 were sequenced from a transformation-ready full-length cDNA library [29], 2851 were collected from NCBI [30]. The EST were annotated against Nr, KEGG, Trembl, Swissprot InterPro and GO databases. The successfully annotated EST sequences were deposited under each database classification.

### Germplasm data collection

Besides the nucleotide data, the HeveaDB database also stored rubber tree germplasm data. Totally 5,049 IRRDB 1981’ wild germplasm accession phenotypic data and 18,328 Wickham rubber clone’s information were deposited. IRRDB 1981’ wild germplasm data was collected from the National Germplasm Garden of Rubber Tree (Danzhou, Hainan) [27], while the Wickham rubber clone’s information was collected from the internal circulated publications and publicly published literatures elsewhere.

### Gene expression analysis

Ninety-nine transcriptomes were mapped to reference genome (Reyan7-33-97) with tophat2 [31], and the FPKM values of each gene were extracted. The 99 transcriptomes belong to 74 samples, some of which have 2 or 3 biological replicates (Supplemental Table 1). The expression value for the samples with biological replicates were calculated from the means, while the value for the samples without replicate was taken from the transcriptome directly. A gene expression matrix was then obtained and used for expression display and co-expression network construction.

### Co-expression network construction

A WGCNA (weighted gene co-expression network analysis) co-expression network [32] was constructed from gene expression data of 74 samples used in expression visualization module. The Pearson’s coefficient threshold value was set as >0.3 to build the network. Some of the genes have too many near nodes, which make the network too complicated to be recognized. We therefore only display the top 20 nodes in the co-expression network. The information of target node genes is also displayed below the network.

### Database construction

The HeveaDB database are currently running on a Linux (CentOS release 6.6) virtual cluster platform. Eight-core processors with 32 GB memory resources were allocated for HeveaDB database. The software environments are set up as the following: Apache Tomcat 7.0.64, Java openjdk-1.8.0.161, Perl 5.10.1, Apache HTTP server 2.2.15, MySQL 5.6.38. Genome were browsed in GBrowse 2.54 and JBrowse 1.12.3. The gene expression is visualized by ECharts 3.2.2, and the co-expression network is drawn with Cytoscape.js 2.7.2. BLAST 2.2.26 and BLAT 3.5 are used to align with the genome. Other pages are generated as static web pages.

## Results and Disccussion

### An intergrated genomic database for rubber tree study

The HeveaDB database is highlighted in four features: 1, The HeveaDB collected substantial data of the rubber trees, including 4 versions of draft genome, 99 NGS transcriptomes, 18 451annotated EST sequences, 12 gene families, 30 200 gene annotations, 5 049 wild germplasm phenotype data, and 18 328 rubber clone’s information etc. (**Table 1**); 2, The HeveaDB provide user-friendly interface to facilitate scientists to utilize the data. Several tools are integrated in the database, including genome browser, blast, blat, text search, sequence retrieve and download etc.; 3, The HeveaDB mined the raw data and provide easy-to-use info to the user. The transcriptomes were analyzed and the gene expression are presented as heatmap and digital FPKM formats. A co-expression network was constructed to show possible correlation between genes; 4, The HeveaDB has a mechanism to receive and share data from rubber tree community under the framework of IRRDB.

**Table 1.**
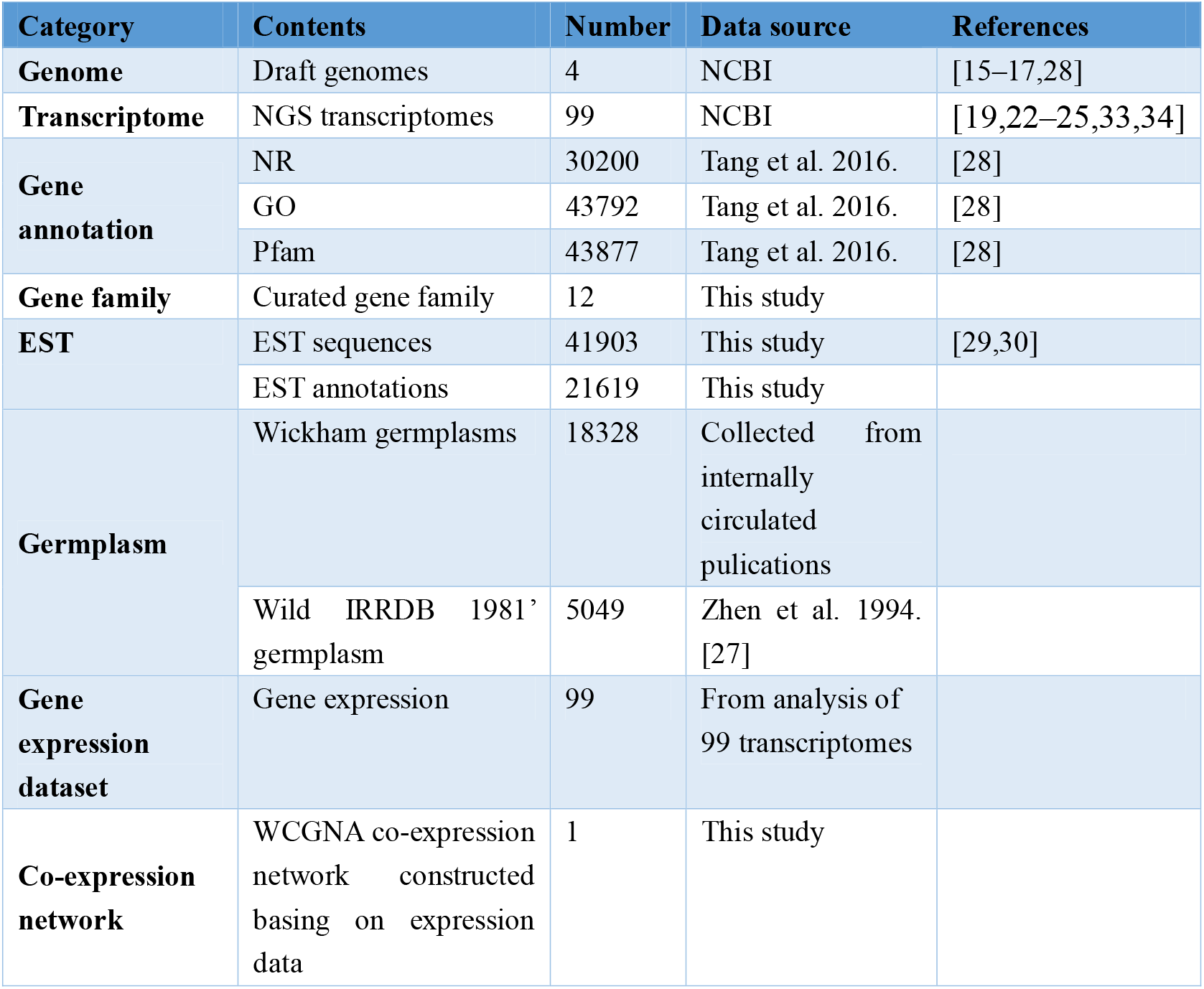
Data content and sources of each database in HeveaDB version 1.0.

| Category | Contents | Number | Data source | References |
| --- | --- | --- | --- | --- |
| **Genome** | Draft genomes | 4 | NCBI | [15–17,28] |
| **Transcriptome** | NGS transcriptomes | 99 | NCBI | [19,22–25,33,34] |
| **Gene annotation** | NR | 30200 | Tang et al. 2016. | [28] |
|  | GO | 43792 | Tang et al. 2016. | [28] |
|  | Pfam | 43877 | Tang et al. 2016. | [28] |
| **Gene family** | Curated gene family | 12 | This study |  |
| **EST** | EST sequences | 41903 | This study | [29,30] |
|  | EST annotations | 21619 | This study |  |
| **Germplasm** | Wickham germplasms | 18328 | Collected from internally circulated publications |  |
|  | Wild IRRDB 1981' germplasm | 5049 | Zhen et al. 1994. [27] |  |
| **Gene expression dataset** | Gene expression | 99 | From analysis of 99 transcriptomes |  |
| **Co-expression network** | WCGNA co-expression network constructed basing on expression data | 1 | This study |  |

### Database structure and organization

The database has 5 major functional modules: Data, Search, Tools, Download and Submit (**Figure 1**). To enable the researcher to explore the rubber tree genome data in a friendly interface, a drop-down menu was provided to access the functions at the upper part of the homepage. A quick search box is presented in the top-right of the page for searching the genes and germplasms.

**Figure 1.**
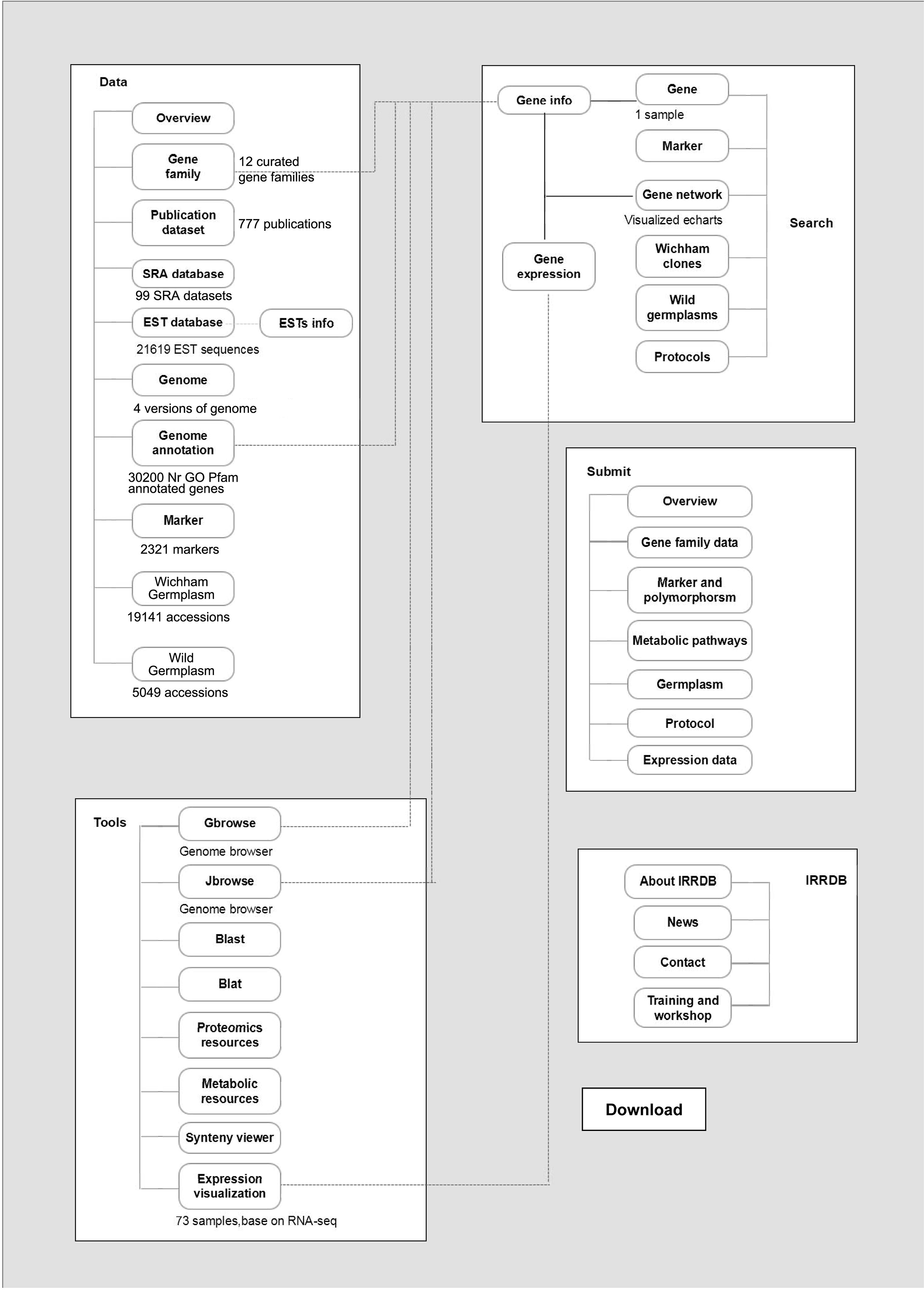
The structure of HeveaDB database. There are 5 major functional modules in the database: Data, Search, Tools, Download and Submit. Each module is connected with gene ID and integrated into gene info pages. The data content and functions are annotated under the block in each module.

### Gene page

Detailed info of each gene is displayed in a compiled gene page. The info includes: ID, loci, structure, expression heatmap, co-expression network, sequences (genomic sequence, transcripts, CDS and deduced peptide), gene annotations (Figure 2). Besides the sequence of CDS and deduced peptide, the up- and down-stream sequences (promoter, terminator etc.) can also be retrieved by entering the numbers to be shown in the genomic sequence panel. The gene pages can be accessed through hyperlinks elsewhere in the HeveaDB database.

**Figure 2.**
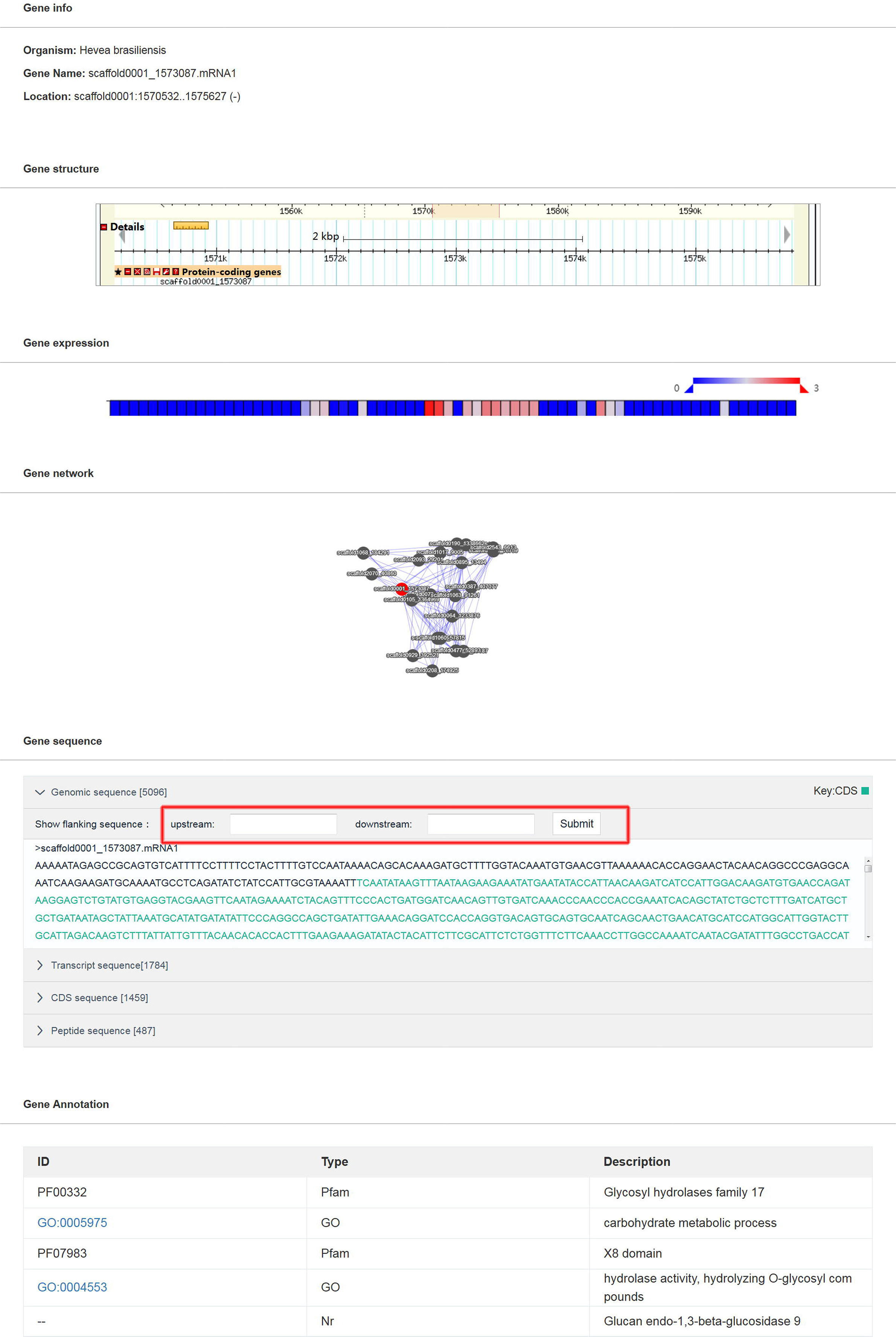
The organization of the gene page. The gene page includes: ID, loci, structure, expression heatmap, co-expression network, sequences (genomic sequence, transcripts, CDS and deduced peptide), gene annotations. The user can retrieve the up- and down-stream sequences of the gene by entering the numbers to be shown in red box.

### Genome browser

The reference genome from CATAS7-33-97 clone is visualized in the Gbrowse and Jbrowse respectively (Figure 3). Each gene is linked to its detail information page by clicking the gene names in the browser (Figure 2). The Gbrowse detail page includes the following basic information of this gene: name, position in the scaffold, length, CDS parts, and sequence. The Jbrowse detail page is showed in a popup window. The primary data and attributes of a gene are displayed, including name, position, length, and region sequences and their sub-features. The default scaffold in the genome browsers is scaffold0001, the user can shift to view particular scaffold by entering the name in the searching form.

**Figure 3.**
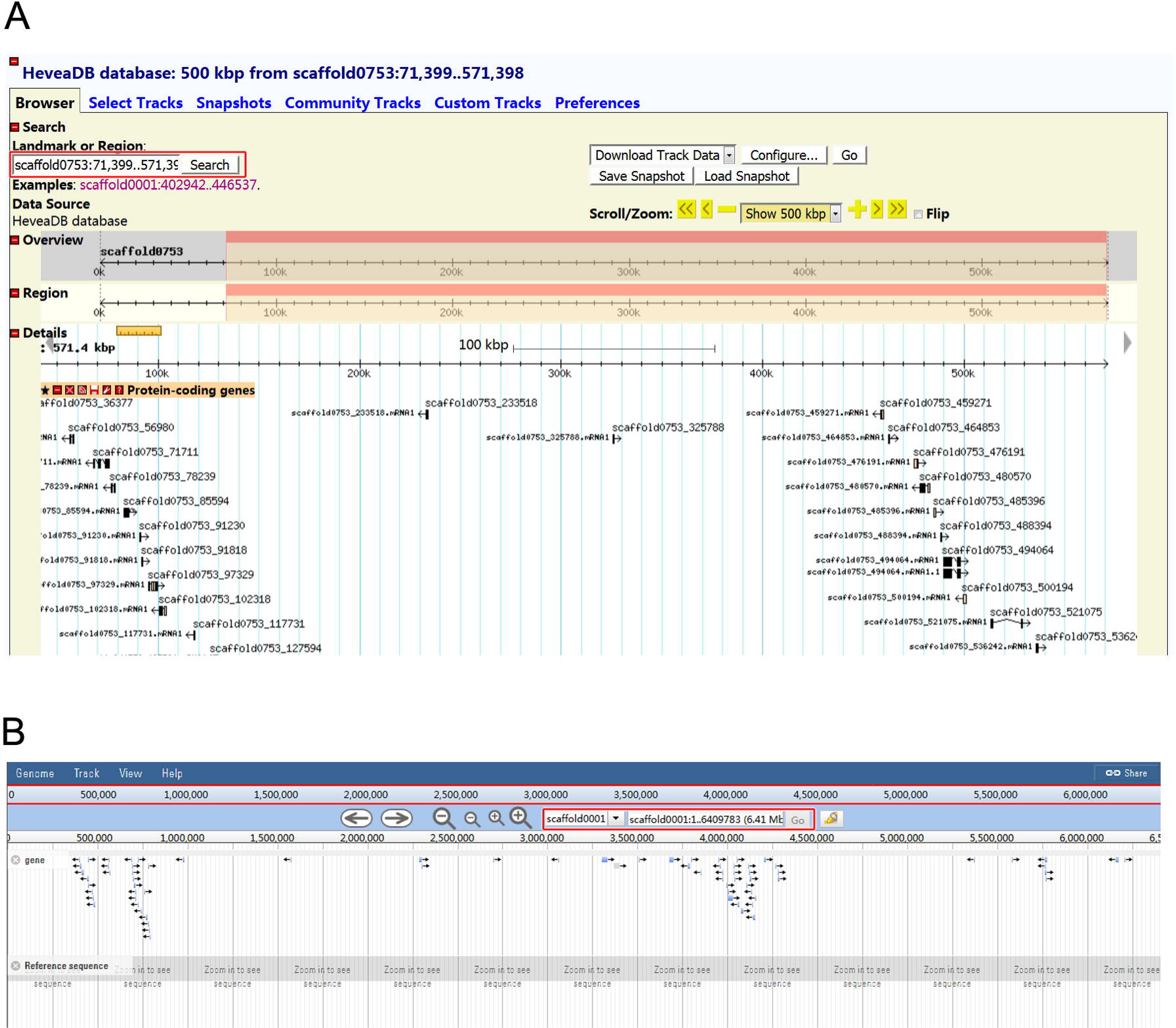
Genome scaffolds are displayed in genome browsers Gbrowse (A) and Jbrowse (B).

### Data search systems

Six types of searching system were provided in HeveaDB (Table 2): gene search, marker search, germplasm search, gene expression search, and sequence homology search (blast and blat). Gene search utilized ID or annotation as keywords to search the functional annotated genes in rubber tree. The genes can be viewed in a compiled gene page, which includes the general gene information, genes structure, gene expression, co-expression network, gene sequences (including CDS and deduced peptide), and annotation (Figure 4). The gene structure was extracted from genome annotation and viewed in the Gbrowse, while the expression of this gene in 74 samples is displayed as heatmap. The rubber genetic map can also be searched by linkage group or marker name (Figure 5). Since there are limited markers in HeveaDB, the markers can also be browsed by clicking the linkage group names at the left side. In gene expression searching system, up-to 20 gene ID can be investigated once. Users can also narrow down the sample numbers through the check boxes which are arranged by clones, tissues and treatments. The expression results are displayed as heatmaps or FPKM values according to the user’s needs (Figure 6). The blast and blat programs were incorporated to provide homology search against genome, CDS and protein sequences (Figure 7). Finally, the information of the Wickham and IRRDB1981 wild germplasm can also be retrieved by search the clone names or germplasm accession (Figure 8). For the Wickham clones, the breeding organism, country, parentages and obtention year are displayed, while for the wild germplasms, both the quantitative (two types of latex yields) and qualitative traits (Girth increment, pulvinus groove, petiolule length, nectary shape, leaf shape, leaf margin, leaf base, leaflets separation, latex color, sprouting stage, defoliation stage and latex yield evaluation) data are displayed (Figure 8).

**Table 2.** Search engines embedded in HeveaDB database.

| **Search Engine** | **Database** | **Keyword type** | **Address** |
| --- | --- | --- | --- |
| **Gene Search** | Gene annotation | Gene ID;<br>Protein ID;<br>Annotation | http://hevea.catas.cn/search/v1/toGeneSearch |
| **Marker Search** | Genetic map | Marker name: | http://hevea.catas.cn/search/v1/toMarker |
| **Network Search** | Co-expression network | Gene ID | http://hevea.catas.cn/coexpression/v1/toCoexpression |
| **Gene Expression Search** | Gene expression database | Gene ID;<br>Sample names | http://hevea.catas.cn/tool/v1/toExpression |
| **Sequence homology search** | Genome database (blast, blat) | Query sequence | http://hevea.catas.cn/tool/v1/toBlat;<br>http://hevea.catas.cn/tool/v1/toBlast |
| **Germplasm Search** | Wickham germplasm;<br>Wild germplasm | Clone accession | http://hevea.catas.cn/search/v1/toGermplasms;<br>http://hevea.catas.cn/search/v1/toPhenotype. |

**Figure 4.**
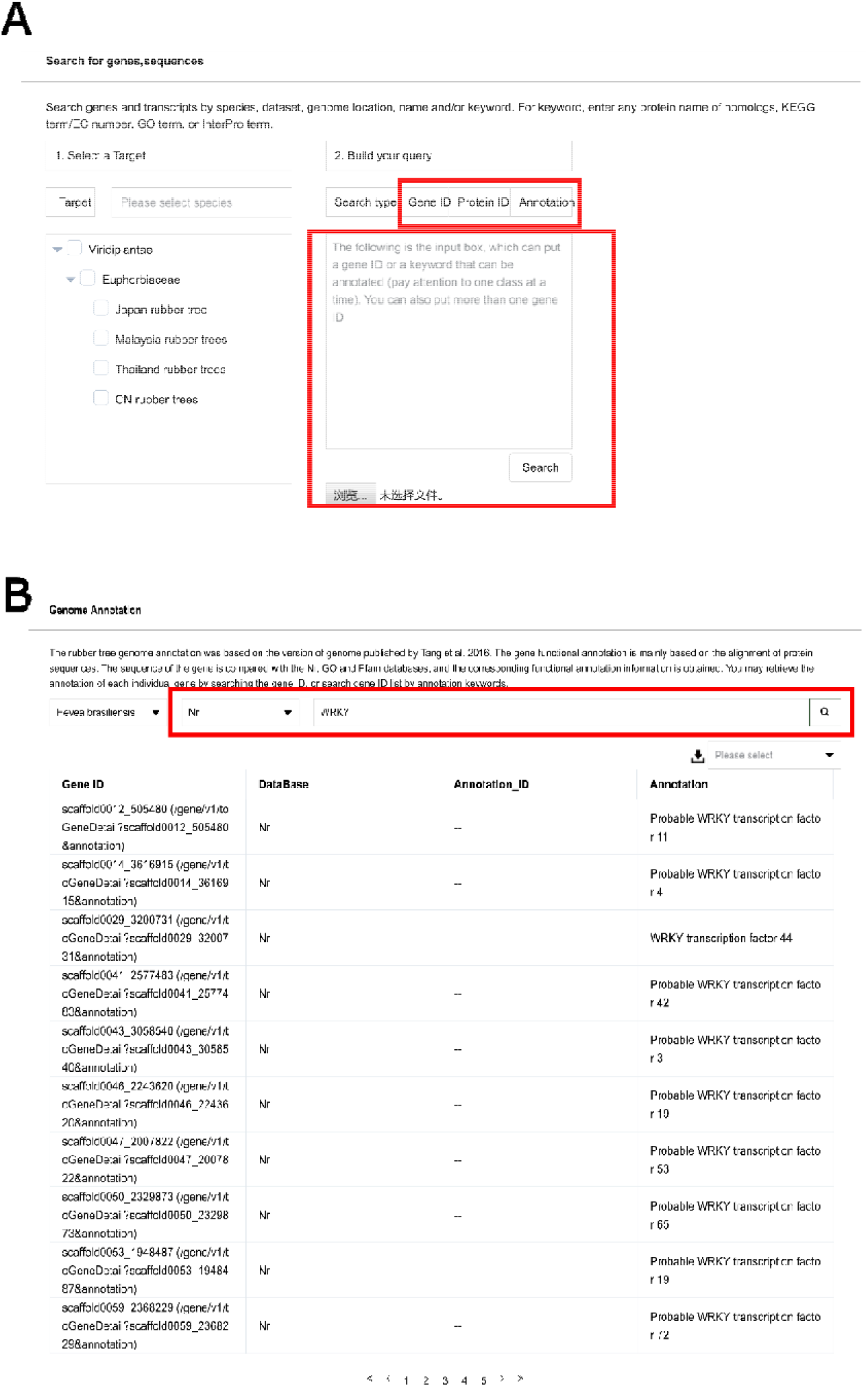
Gene searching engine (A) and multiple genes returned through annotation keywords search (B). Gene ID, Protein ID and annotation text can be used as keywords to search the functional annotated genes in rubber tree. When annotation text is used as keywords, multiple hits will be returned as a list.

**Figure 5.**
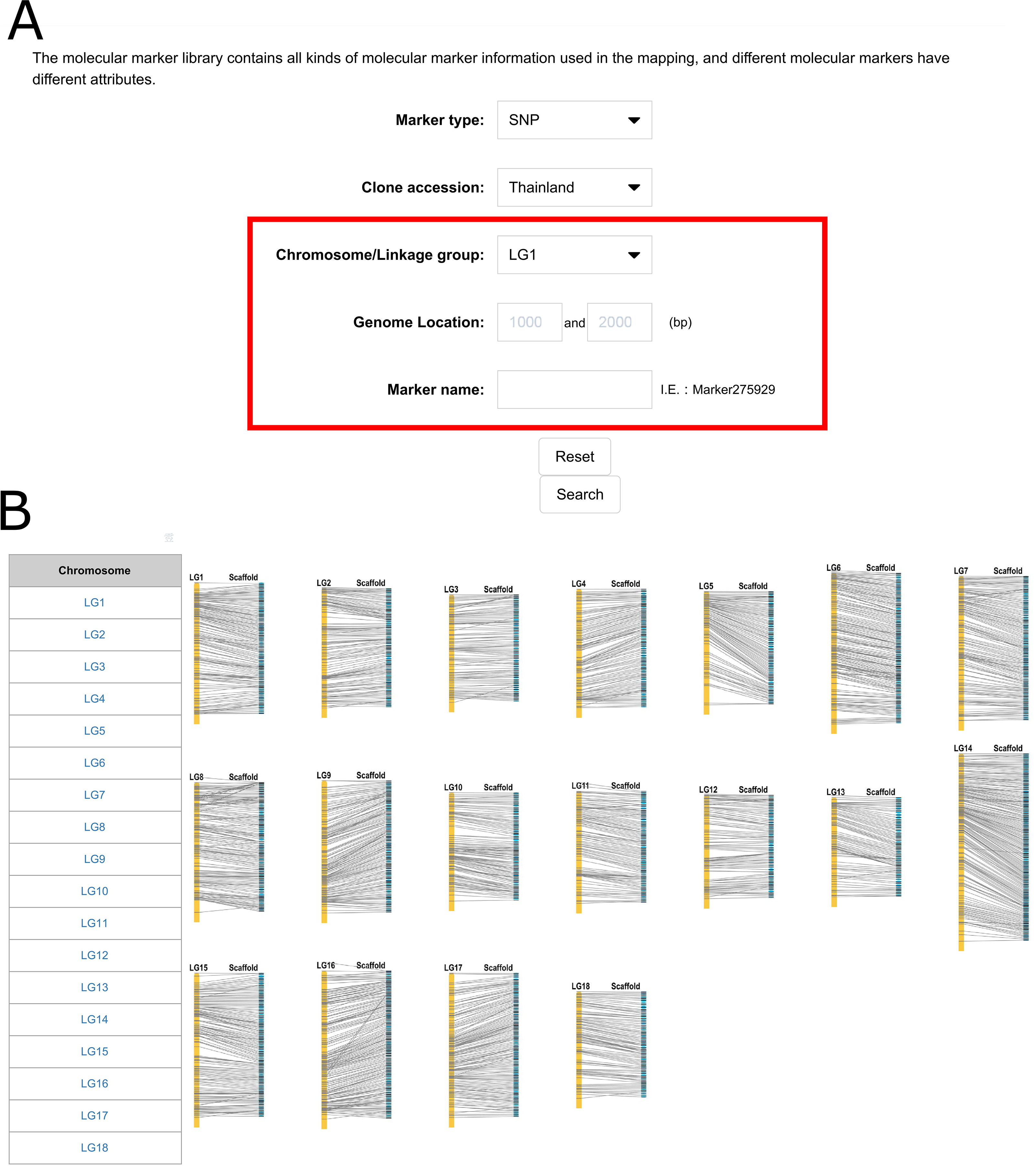
Molecular marker searching system (A) and a genetic map with 18 linkage groups to be displayed by clicking (B).

**Figure 6.**
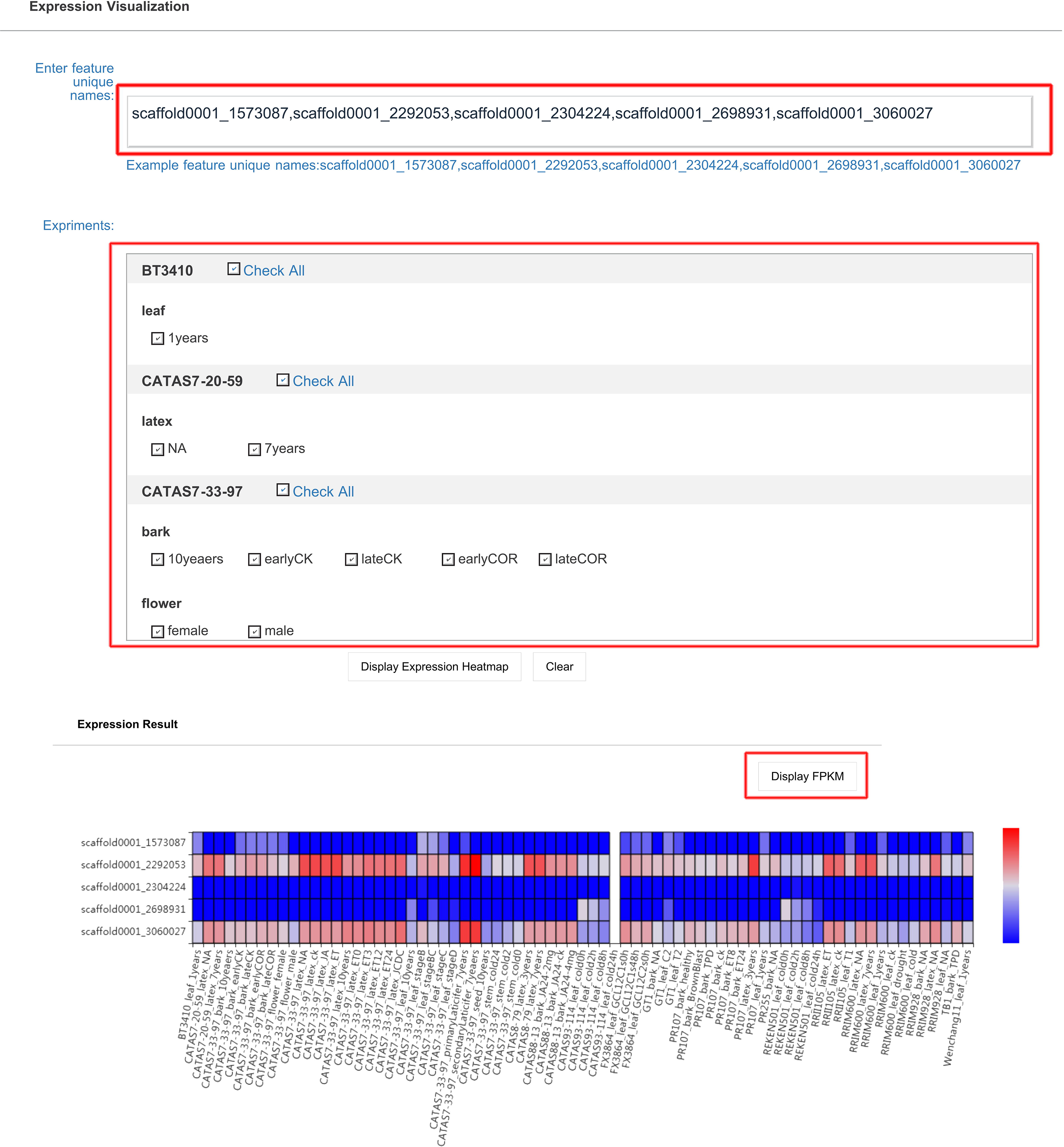
Gene expression searching and visualization system. Up-to 20 gene ID can be investigated once. Users can also narrow down the sample numbers through the check boxes which are arranged by clones, tissues and treatments. The expression results are displayed as heatmaps or FPKM values according to the user’s needs. Each row represents the expression data of a gene, and each column shows the expression in a sample.

**Figure 7.**
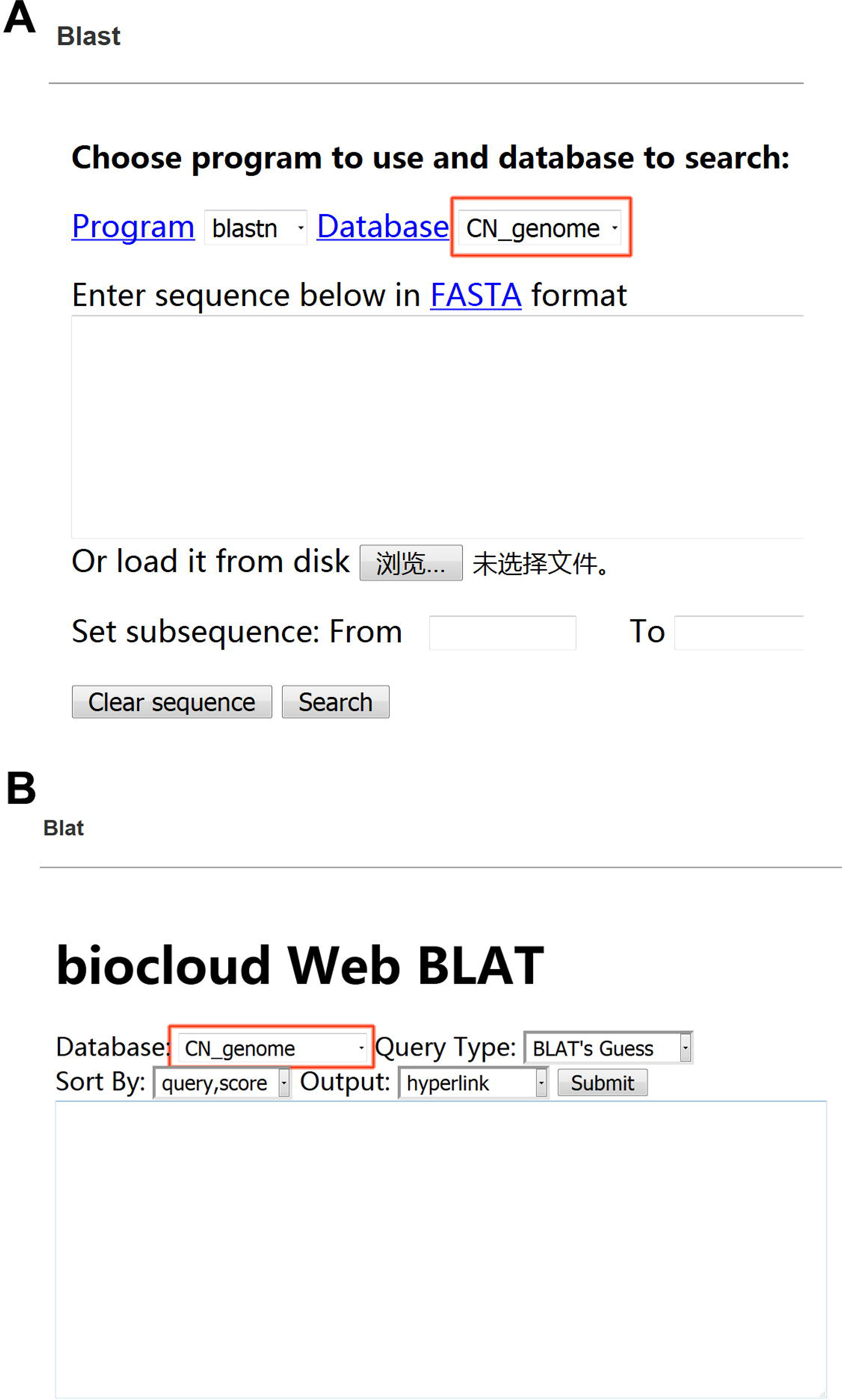
The integrated Blast (A) and Blat (B) programs to search against genome, CDS and protein sequences.

**Figure 8.**
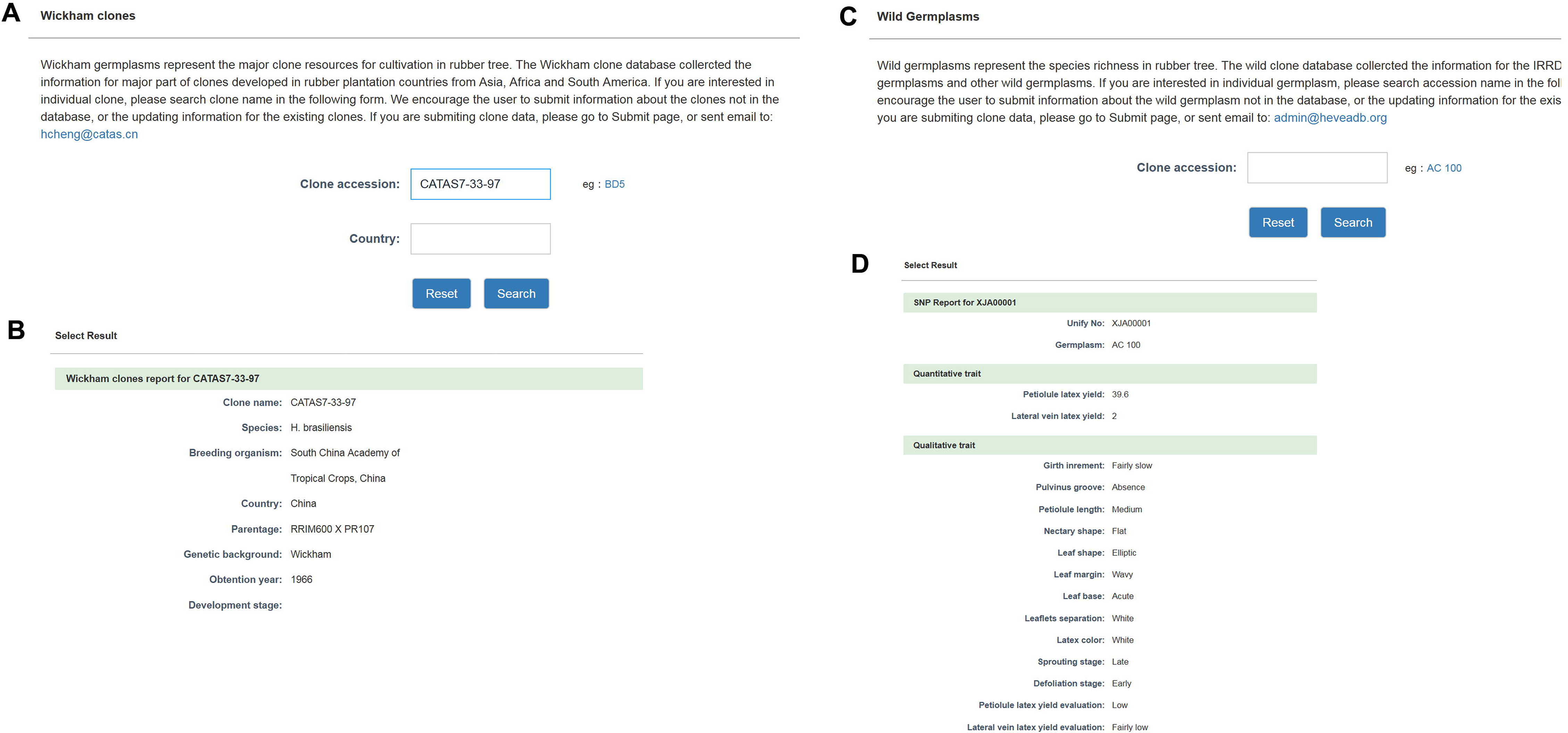
The germplasm searching systems. A, B, the Wickham clone search system and the information provided. C, D, the IRRDB1981 wild germplasm searching system and the information provided.

### Other pages

Other data and information, including data submission instructions, are also incorporated in the HeveaDB database. IRRDB information pages, which provide basic descriptions about this database and IRRDB community.

## Perspectives and concluding remarks

The fast proceeding of sequencing technology greatly benefits the genome study in non-model plant species, such as *Hevea brasiliensis*. The appearance of huge amount of dataset requires a comprehensive database for the researchers in rubber tree community. The HeveaDB is a datahub to store, share, distribute and re-utilize rubber tree NGS data. Using CATAS7-33-97 genome as the reference, HeveaDB provide the genome visualization, gene annotation, co-expression network, and expression visualization. Besides, the data is also searchable through the searching module, and downloadable through the web engine. Though HeveaDB is still a preliminary genomic database system, it has achieved much attention and usage in rubber community. Until now, HeveaDB achieved 8,499 pageviews from 1491 unique viewers of 14 countries and districts in the past one year (Supplemental Table 2). To make HeveaDB easier to access and more powerful to employ, further improvement will be conducted. With the progress on rubber tree genome study, the data will be updated periodically. More functions will be developed to facilitate post-genomic study analysis online. Now the rubber community has began to implement the resequencing project of Hevea germplasm. The HeveaDB will focus on the data collection of both genomic and phenomic data, and to establish a platform for the potential new discoveries for genomic studies and breeding technology in rubber tree.

## Author contributions

CH, XY and HH designed the database, CH collected the data and calculated the gene expression from transcriptomes. ZJ coordinated the cooperation between IRRDB members, CH wrote the manuscript and HH revised.

## Competing interests

The authors declare that the research was conducted in the absence of any commercial or financial relationships that could be construed as a potential conflict of interest.

## Acknowledgements

The HeveaDB datahub project was led by the IRRDB Molecular Biology and Physiology (MBP) and was financially supported by the Central Public-interest Scientific Institution Basal Research Fund for CATAS (No. 1630022018007) and the Fund for Innovative Research Team Program of CATAS (NO. 17CXTD-28). I thank Dr. Chow Keng See from Malaysian Rubber Board, Dr. Fang Yongjun and Dr. An Zewei from CATAS for their insightful suggestions for the construction of HeveaDB database.

## Notes

http://hevea.catas.cn

