## Supplemental Table 1 for "HeveaDB: a genetic resource database for rubber tree genomic study"

Table 3 Ninty-nine selected transcriptomes used for gene expression analysis.

| Transcriptome | Duplicates | SRA accession No. | Sample name | Description |
| --- | --- | --- | --- | --- |
| H1 | 2 | SRR854520 | RRIM600_latex_- | Latex samples from the RRIM600 clone |
| H2 | 2 | SRR854521 | RRIM600_latex_- | Latex samples from the RRIM600 clone |
| H3 | 2 | SRR854522 | CATAS7-20-59_latex_- | Latex samples from the CATAS7-20-59 clone. |
| H4 | 2 | SRR1533844 | CATAS7-20-59_latex_- | Latex samples from the CATAS7-20-59 clone. |
| H5 | 1 | SRR1533845 | PR107_latex_3years | Latex samples from 3 years old PR107 clone. |
| H6 | 1 | SRR1544255 | CATAS8-79_latex_3years | Latex samples from 3 years old CATAS8-79 clone. |
| H7 | 1 | SRR1544256 | PR255_bark_- | Bark samples from PR255 clone. |
| H8 | 1 | SRR1588170 | GT1_bark_- | Bark samples from GT1 clone. |
| H9 | 2 | SRR1611790 | PR107_bark_healthy | Bark samples from 14 years old healthy PR107 clone |
| H10 | 2 | SRR1611791 | PR107_bark_healthy | Bark samples from 14 years old healthy PR107 clone |
| H11 | 1 | SRR1611792 | PR107_bark_BrownBlast | Bark samples from 14 years old PR107 clone with brown blast disease |
| H12 | 1 | SRR1648124 | PR107_bark_TPD | Bark samples from 14 years old PR10 clone with TPD disease |
| H13 | 1 | SRR167696 | CATAS7-33-97_latex_- | Latex samples from the CATAS7-33-97 clone. |
| H14 | 1 | SRR2001618 | CATAS7-33-97_bark-leaf | Bark and leaf mixture samples from the CATAS7-33-97 clone |
| H15 | 1 | SRR2001619 | RRIM600_latex_7years | Latex samples from 7 years old RRIM600 clone. |
| H16 | 1 | SRR2001620 | CATAS7-20-59_latex_7years | Latex samples from 7 years old CATAS7-20-59 clone. |
| H17 | 1 | SRR2002800 | CATAS8-79_latex_7years | Latex samples from 7 years old CATAS8-79 clone. |
| H18 | 1 | SRR2002803 | CATAS7-33-97_latex_ck | Latex samples from 7 years old CATAS7-33-97 clone as control. |
| H19 | 1 | SRR2002804 | CATAS7-33-97_latex_JA | Latex samples from 7 years old CATAS7-33-97 clone treated with JA-2 |
| H20 | 1 | SRR2063953 | CATAS7-33-97_latex_ET | Latex samples from 7 years old CATAS7-33-97 clone treated with ET-2. |
| H21 | 2 | SRR2063959 | TB1_bark_TPD | Bark samples from TB1 in TPD. |
| H22 | 2 | SRR2147072 | TB1_bark_TPD | Bark samples from TB1 in TPD. |
| H23 | 1 | SRR2147073 | PR107_bark_ck | Bark samples from 15 years old PR107 clone as control |
| H24 | 1 | SRR2147074 | PR107_bark_ET8 | Bark samples from 15 years old PR107 clone treated with 1.5% Ethephon for 8 hours |
| H25 | 1 | SRR2156988 | PR107_bark_ET24 | Bark samples from 15 years old PR107 clone treated with 1.5% Ethephon for 24 hours |
| H26 | 1 | SRR2156992 | FX3864_leaf_GCL12C1s0h | Leaf samples from FX3864 clone traeted with GCL12C1s for 0h |
| H27 | 1 | SRR2157179 | FX3864_leaf_GCL12C1s48h | Leaf samples from FX3864 clone traeted with GCL12C1s for 48h |
| H28 | 1 | SRR3136158 | FX3864_leaf_GCL12C2s0h | Leaf samples from FX3864 clone traeted with GCL12Cs for 0h |
| H29 | 1 | SRR3136159 | CATAS7-33-97_bark_10yeaers | Bark samples from 10 years old CATAS7-33-97 clone. |
| H30 | 1 | SRR3136162 | CATAS7-33-97_leaf_10years | Leaf samples from 10 years old CATAS7-33-97 clone. |
| H31 | 1 | SRR3136165 | CATAS7-33-97_latex_10years | Latex samples from 10 years old CATAS7-33-97 clone. |
| H32 | 1 | SRR3136166 | CATAS7-33-97_flower_female | Femail flower samples from 10 years old CATAS7-33-97 clone. |
| H33 | 1 | SRR3136168 | CATAS7-33-97_flower_male | Male flower samples from 10 years old CATAS7-33-97 clone. |
| H34 | 1 | SRR3136173 | CATAS7-33-97_seed_10years | Seeds samples from 10 years old CATAS7-33-97 clone. |
| H35 | 1 | SRR3136176 | CATAS7-33-97_latex_ET0 | Latex samples from 10 years old CATAS7-33-97 clone treated with Ethrel for 0h as control. |
| H36 | 1 | SRR3136177 | CATAS7-33-97_latex_ET3 | Latex samples from 10 years old CATAS7-33-97 clone treated with Ethrel for 3h. |
| H37 | 1 | SRR3136178 | CATAS7-33-97_latex_ET12 | Latex samples from 10 years old CATAS7-33-97 clone treated with Ethrel for 12h. |
| H38 | 1 | SRR3136185 | CATAS7-33-97_latex_ET24 | Latex samples from 10 years old CATAS7-33-97 clone treated with Ethrel for 24h. |
| H39 | 1 | SRR3136188 | CATAS7-33-97_leaf_stageB | Leaf samples in stage B from 10 years old CATAS7-33-97 clone. |
| H40 | 1 | SRR3136190 | CATAS7-33-97_leaf_stageBC | Leaf samples in stage B-C from 10 years old CATAS7-33-97 clone. |
| H41 | 1 | SRR3136192 | CATAS7-33-97_leaf_stageC | Leaf samples in stage C from 10 years old CATAS7-33-97 clone. |
| H42 | 1 | SRR3240371 | CATAS7-33-97_leaf_stageD | Leaf samples in stage D from 10 years old CATAS7-33-97 clone. |
| H43 | 1 | SRR3240372 | PR107_leaf_1years | Leaf samples from 1 years old PR107 clone |
| H44 | 1 | SRR3240373 | RRIM600_leaf_1years | Leaf samples from 1 years old RRIM600 clone |
| H45 | 1 | SRR3240374 | BT3410_leaf_1years | Leaf samples from 1 years old BT3410 clone |
| H46 | 1 | SRR3423347 | Wenchang11_leaf_1years | Leaf samples from 1 years old Wenchang11 clone |
| H47 | 1 | SRR3423348 | CATAS7-33-97_bark_earlyCK | Bark samples from CATAS7-33-97 clone as early control |
| H48 | 1 | SRR3423349 | CATAS7-33-97_bark_lateCK | Bark samples from CATAS7-33-97 clone as late control |
| H49 | 1 | SRR3423350 | CATAS7-33-97_bark_earlyCoronatine | Bark samples from CATAS7-33-97 clone treated with Coronatine at early stage. |
| H50 | 1 | SRR5051416 | CATAS7-33-97_bark_lateCoronatine | Bark samples from CATAS7-33-97 clone treated with Coronatine at late stage. |
| H51 | 1 | SRR5051417 | CATAS88-13_bark_JA24-2mg | Bark samples from CATAS88-13 clone treated with 2mg JA for 24h. |
| H52 | 1 | SRR5051418 | CATAS88-13_bark_JA24-ck | Bark samples from CATAS88-13 clone as ck for 24h. |
| H53 | 1 | SRR5118395 | CATAS88-13_bark_JA24-4mg | Bark samples from CATAS88-13 clone treated with 4mg JA for 24h. |
| H54 | 3 | SRR5118396 | CATAS7-33-97_secondaryLaticifer_7yeaers | Secondary laticifer samples from 7 years old CATAS7-33-97 clone |
| H55 | 3 | SRR5118397 | CATAS7-33-97_primaryLaticifer_7years | Primary laticifer samples from 7 years old CATAS7-33-97 clone |
| H56 | 3 | SRR5118398 | CATAS7-33-97_secondaryLaticifer_7yeaers | Secondary laticifer samples from 7 years old CATAS7-33-97 clone |
| H57 | 3 | SRR5118399 | CATAS7-33-97_primaryLaticifer_7years | Primary laticifer samples from 7 years old CATAS7-33-97 clone |
| H58 | 3 | SRR5118400 | CATAS7-33-97_secondaryLaticifer_7yeaers | Secondary laticifer samples from 7 years old CATAS7-33-97 clone |
| H59 | 3 | SRR5868395 | CATAS7-33-97_primaryLaticifer_7years | Primary laticifer samples from 7 years old CATAS7-33-97 clone |
| H60 | 2 | SRR5868396 | CATAS7-33-97_latex_JCDC | Latex samples from 3 years old CATAS7-33-97 clone |
| H61 | 2 | SRR611643 | CATAS7-33-97_latex_JCDC | Latex samples from 3 years old CATAS7-33-97 clone |
| H62 | 1 | SRR611644 | GT1_leaf_C2 | Leaf samples from GT1 clone with Corynespora cassiicola tolerance as control |
| H63 | 1 | SRR611645 | RRII105_leaf_T1 | Leaf samples from RRII105 clone with Corynespora cassiicola tolerance as T1 treatment |
| H64 | 1 | SRR6127583 | GT1_leaf_T2 | Leaf samples from GT1 clone with Corynespora cassiicola tolerance as T2 treatment |
| H65 | 1 | SRR6127584 | CATAS7-33-97_stem_cold24 | Stem samples from 1 years old CATAS7-33-97 clone treated under 4 C for 24 h |
| H66 | 1 | SRR6127585 | CATAS7-33-97_stem_cold2 | Stem samples from 1 years old CATAS7-33-97 clone treated under 4 C for 2 h |
| H67 | 1 | SRR620233 | CATAS7-33-97_stem_cold0 | Stem samples from 1 years old CATAS7-33-97 clone treated under 4 C for 0 h |
| H68 | 1 | SRR620234 | RRIM600_leaf_ck | Leaf samples from 6 months old RRIM600 clone as control |
| H69 | 1 | SRR620235 | RRIM600_leaf_drought | Leaf samples from 6 months old RRIM600 clone treated with drought |
| H70 | 1 | SRR620236 | RRIM600_leaf_cold | Leaf samples from 6 months old RRIM600 clone treated with cold stress. |
| H71 | 1 | SRR620237 | RRII105_latex_ET | Latex samples from 20 years old RRII105 clones treated with ethylene |
| H72 | 1 | SRR1508164 | RRII105_latex_ck | Latex samples from 20 years old RRII105 clones as control |
| H73 | 1 | SRR1508165 | RRIM928_bark_- | Bark samples from the RRIM928 clone |
| H74 | 1 | SRR1508166 | RRIM928_latex_- | Latex samples from the RRIM928 clone |
| H75 | 1 | SRR1508167 | RRIM928_leaf_- | Leaf samples from the RRIM928 clone |
| H76 | 3 | SRR6668812 | REKEN501_leaf_cold0h | Leaf samples from 2 years old Reken501 clone treated with cold for 0h |
| H77 | 3 | SRR6668813 | REKEN501_leaf_cold0h | Leaf samples from 2 years old Reken501 clone treated with cold for 0h |
| H78 | 3 | SRR6668814 | REKEN501_leaf_cold0h | Leaf samples from 2 years old Reken501 clone treated with cold for 0h |
| H79 | 3 | SRR6668811 | REKEN501_leaf_cold24h | Leaf samples from 2 years old Reken501 clone treated with cold for 24h |
| H80 | 3 | SRR6668806 | REKEN501_leaf_cold24h | Leaf samples from 2 years old Reken501 clone treated with cold for 24h |
| H81 | 3 | SRR6668807 | REKEN501_leaf_cold24h | Leaf samples from 2 years old Reken501 clone treated with cold for 24h |
| H82 | 3 | SRR6668815 | REKEN501_leaf_cold2h | Leaf samples from 2 years old Reken501 clone treated with cold for 2h |
| H83 | 3 | SRR6668816 | REKEN501_leaf_cold2h | Leaf samples from 2 years old Reken501 clone treated with cold for 2h |
| H84 | 3 | SRR6668817 | REKEN501_leaf_cold2h | Leaf samples from 2 years old Reken501 clone treated with cold for 2h |
| H85 | 3 | SRR6668818 | REKEN501_leaf_cold8h | Leaf samples from 2 years old Reken501 clone treated with cold for 8h |
| H86 | 3 | SRR6668819 | REKEN501_leaf_cold8h | Leaf samples from 2 years old Reken501 clone treated with cold for 8h |
| H87 | 3 | SRR6668810 | REKEN501_leaf_cold8h | Leaf samples from 2 years old Reken501 clone treated with cold for 8h |
| H88 | 3 | SRR6668820 | CATAS93-114_leaf_cold24h | Leaf samples from 2 years old CATAS93-114 clone treated with cold for 24h |
| H89 | 3 | SRR6668823 | CATAS93-114_leaf_cold24h | Leaf samples from 2 years old CATAS93-114 clone treated with cold for 24h |
| H90 | 3 | SRR6668822 | CATAS93-114_leaf_cold24h | Leaf samples from 2 years old CATAS93-114 clone treated with cold for 24h |
| H91 | 3 | SRR6668808 | CATAS93-114_leaf_cold0h | Leaf samples from 2 years old CATAS93-114 clone treated with cold for 0h |
| H92 | 3 | SRR6668809 | CATAS93-114_leaf_cold0h | Leaf samples from 2 years old CATAS93-114 clone treated with cold for 0h |
| H93 | 3 | SRR6668802 | CATAS93-114_leaf_cold0h | Leaf samples from 2 years old CATAS93-114 clone treated with cold for 0h |
| H94 | 3 | SRR6668803 | CATAS93-114_leaf_cold2h | Leaf samples from 2 years old CATAS93-114 clone treated with cold for 2h |
| H95 | 3 | SRR6668804 | CATAS93-114_leaf_cold2h | Leaf samples from 2 years old CATAS93-114 clone treated with cold for 2h |
| H96 | 3 | SRR6668805 | CATAS93-114_leaf_cold2h | Leaf samples from 2 years old CATAS93-114 clone treated with cold for 2h |
| H97 | 3 | SRR6668800 | CATAS93-114_leaf_cold8h | Leaf samples from 2 years old CATAS93-114 clone treated with cold for 8h |
| H98 | 3 | SRR6668801 | CATAS93-114_leaf_cold8h | Leaf samples from 2 years old CATAS93-114 clone treated with cold for 8h |
| H99 | 3 | SRR6668821 | CATAS93-114_leaf_cold8h | Leaf samples from 2 years old CATAS93-114 clone treated with cold for 8h |
