## Supplementary material for "HeveaDB: a genetic resource database for rubber tree genomic study": Supplmental Table 2

Supplemental Table 2. The usage data of HeveaDB database (2018.4.27~2019.4.3)

|  | Page Views | Unique visitors | IP | Visit View | PV Per User Per Visit | Depth of Visit |
| --- | --- | --- | --- | --- | --- | --- |
| China | 8108 | 1393 | 1061 | 2278 | 112.19 | 85.25 |
| America | 102 | 12 | 12 | 13 | 59.67 | 47.5 |
| Malaysia | 74 | 16 | 11 | 21 | 4.63 | 3.52 |
| UK | 65 | 46 | 46 | 49 | 1.41 | 1.33 |
| Sri Lanka | 57 | 5 | 4 | 11 | 11.4 | 5.18 |
| Austria | 28 | 3 | 4 | 5 | 9.33 | 5.6 |
| French | 27 | 8 | 8 | 9 | 3.38 | 3 |
| Brasil | 16 | 1 | 1 | 1 | 16 | 16 |
| Vietnam | 10 | 2 | 2 | 2 | 5 | 5 |
| Korea | 4 | 1 | 1 | 1 | 4 | 4 |
| Côte d'Ivoire | 4 | 0 | 1 | 0 | 0 | 0 |
| Thailand | 2 | 2 | 2 | 2 | 1 | 1 |
| Hong Kong | 1 | 1 | 1 | 1 | 1 | 1 |
| Russia | 1 | 1 | 1 | 1 | 1 | 1 |
| Sum | **8499** | **1491** | **1155** | 2394 |  |  |
